## Supplemental Figure 1 for "Metagenomic identification of viral sequences in laboratory reagents"

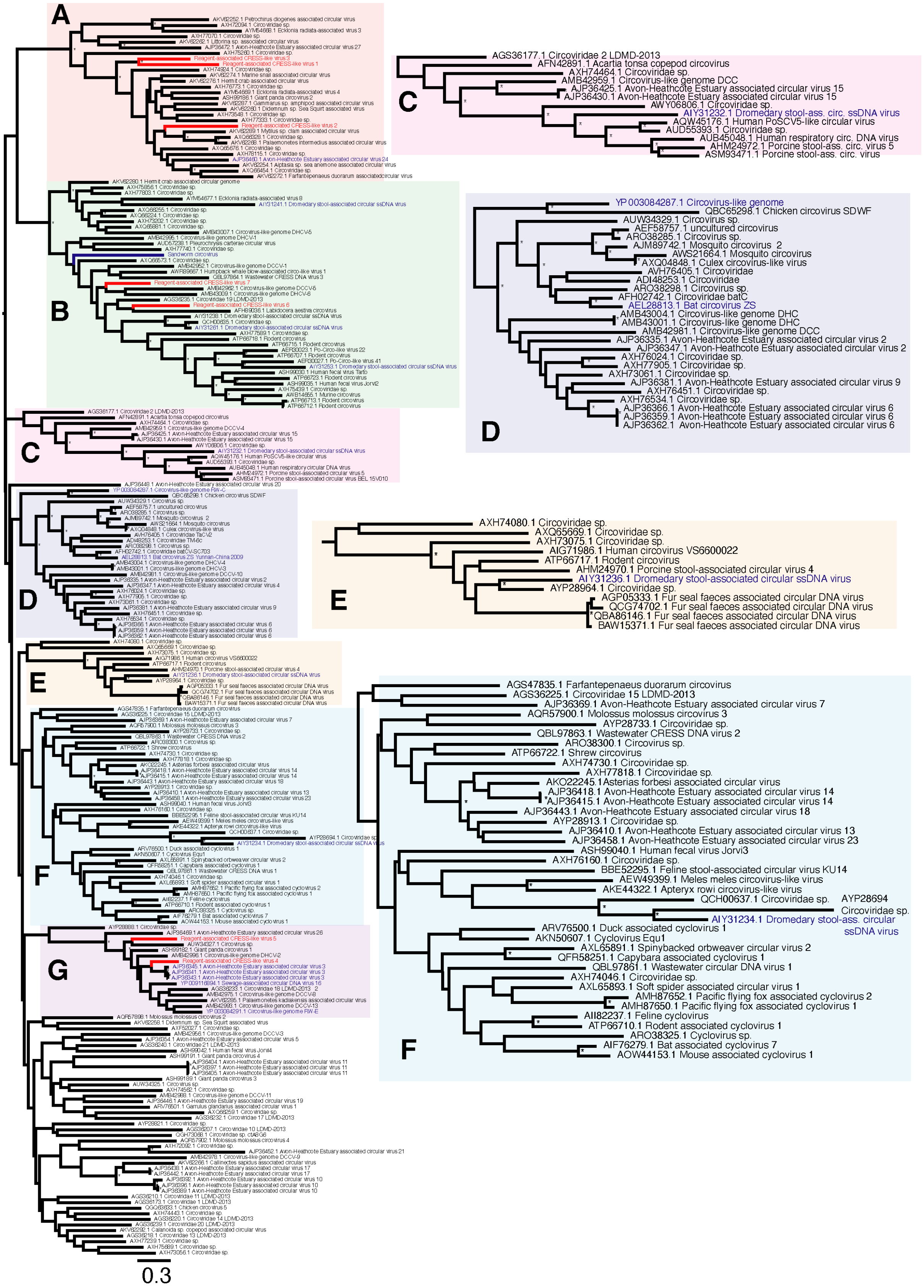
A
B
C
AGS36177.1 Circoviridae 2 LDMD-2013
AFN42891.1 Acartia tonsa copepod circovirus
AXH74464.1 Circoviridae sp.
AMB42959.1 Circovirus-like genome DCC
AJP36425.1 Avon-Heathcote Estuary associated circular virus 15
AJP36430.1 Avon-Heathcote Estuary associated circular virus 15
AWY06806.1 Circoviridae sp.
AIY31232.1 Dromedary stool-ass. circ. ssDNA virus
AQW45176.1 Human PoSCV5-like circular virus
AUD55393.1 Circoviridae sp.
AUB45048.1 Human respiratory circ. DNA virus
AHM24972.1 Porcine stool-ass. circ. virus 5
ASM93471.1 Porcine stool-ass. circ. virus
D
YP 003084287.1 Circovirus-like genome
QBC65298.1 Chicken circovirus SDWF
AUW34329.1 Circovirus sp.
AEF58757.1 uncultured circovirus
ARO38285.1 Circovirus sp.
AJM89742.1 Mosquito circovirus 2
AWS21664.1 Mosquito circovirus
AXQ04848.1 Culex circovirus-like virus
AVH76405.1 Circoviridae
ADI48253.1 Circoviridae
ARO38298.1 Circovirus sp.
AFH02742.1 Circoviridae batC
AEL28813.1 Bat circovirus ZS
AMB43004.1 Circovirus-like genome DHC
AMB43001.1 Circovirus-like genome DHC
AMB42981.1 Circovirus-like genome DCC
AJP36335.1 Avon-Heathcote Estuary associated circular virus 2
AJP36347.1 Avon-Heathcote Estuary associated circular virus 2
AXH76024.1 Circoviridae sp.
AXH77905.1 Circoviridae sp.
AXH73061.1 Circoviridae sp.
AJP36381.1 Avon-Heathcote Estuary associated circular virus 9
AXH76451.1 Circoviridae sp.
AXH76534.1 Circoviridae sp.
AJP36366.1 Avon-Heathcote Estuary associated circular virus 6
AJP36359.1 Avon-Heathcote Estuary associated circular virus 6
AJP36362.1 Avon-Heathcote Estuary associated circular virus 6
E
AXH74080.1 Circoviridae sp.
AXQ65669.1 Circoviridae sp.
AXH73075.1 Circoviridae sp.
AIG71986.1 Human circovirus VS6600022
ATP66717.1 Rodent circovirus
AHM24970.1 Porcine stool-associated circular virus 4
AIY31236.1 Dromedary stool-associated circular ssDNA virus
AYP28964.1 Circoviridae sp.
AGP05333.1 Fur seal faeces associated circular DNA virus
QCG74702.1 Fur seal faeces associated circular DNA virus
QBA86146.1 Fur seal faeces associated circular DNA virus
BAW15371.1 Fur seal faeces associated circular DNA virus
F
AGS47835.1 Farfantepenaeus duorarum circovirus
AGS36225.1 Circoviridae 15 LDMD-2013
AJP36369.1 Avon-Heathcote Estuary associated circular virus 7
AQR57900.1 Molossus molossus circovirus 3
AYP28733.1 Circoviridae sp.
QBL97863.1 Wastewater CRESS DNA virus 2
ARO38300.1 Circovirus sp.
ATP66722.1 Shrew circovirus
AXH74730.1 Circoviridae sp.
AXH77818.1 Circoviridae sp.
AKO22245.1Asterias forbesi associated circular virus
AJP36418.1 Avon-Heathcote Estuary associated circular virus 14
AJP36415.1 Avon-Heathcote Estuary associated circular virus 14
AJP36443.1 Avon-Heathcote Estuary associated circular virus 18
AYP28913.1 Circoviridae sp.
AJP36410.1 Avon-Heathcote Estuary associated circular virus 13
AJP36458.1 Avon-Heathcote Estuary associated circular virus 23
ASH99040.1 Human fecal virus Jorvi3
AXH76160.1 Circoviridae sp.
BBE52295.1 Feline stool-associated circular virus KU14
AEW49399.1 Meles meles circovirus-like virus
AKE44322.1 Apteryx rowi circovirus-like virus
QCH00637.1 Circoviridae sp.
AYP28694 Circoviridae sp.
AIY31234.1 Dromedary stool-ass. circular ssDNA virus
ARV76500.1 Duck associated cyclovirus 1
AKN50607.1 Cyclovirus Equ1
AXL65891.1 Spinybacked orbweaver circular virus 2
QFR58251.1 Capybara associated cyclovirus 1
QBL97861.1 Wastewater circular DNA virus 1
AXH74046.1 Circoviridae sp.
AXL65893.1 Soft spider associated circular virus 1
AMH87652.1 Pacific flying fox associated cyclovirus 2
AMH87650.1 Pacific flying fox associated cyclovirus 1
AII82237.1 Feline cyclovirus
ATP66710.1 Rodent associated cyclovirus 1
ARO38325.1 Cyclovirus sp.
AIF76279.1 Bat associated cyclovirus 7
AOW44153.1 Mouse associated cyclovirus 1
G
0.3

**Supplementary Figure 1.** Phylogenetic relationships of CRESS viruses, including the 7 novel CRESS-like viruses identified in this study. Novel viruses are highlighted in red (Reagent-associated CRESS-like virus 1-7). The clades (C, D, E and F) are shown in higher resolution on the right. For clarity, the tree was mid-point rooted. Bootstrap values greater than 70% are represented by asterisks next to nodes. All horizontal branch lengths are scaled according to number of amino acid substitutions per site.
